# Angiopoietin-like 4 promotes the proliferation and migration of epidermal stem cells and contributes to the re-epithelialization of cutaneous wounds

**DOI:** 10.1101/2023.02.23.529672

**Authors:** Yuan Yang, Chenghao Yu, Yingying Le, Weijuan Gong, Jihui Ju, Guangliang Zhang, Pengxiang Ji, Rui Zuo, Zhe Liu, Ping Zhang, Ruixing Hou, Yi Fu

## Abstract

Proliferation and migration of epidermal stem cells (EpSCs) are essential for epithelialization during skin wound healing. Angiopoietin-like 4 (ANGPTL4) has been reported to play an important role in wound healing, but the mechanisms involved are not fully understood. Here we investigate the contribution of ANGPTL4 to full-thickness wound re-epithelialization and the underlying mechanisms using *Angptl4* knockout mice. Immunohistochemical staining reveals that ANGPTL4 is significantly upregulated in the basal layer cells of the epidermis around the wound during cutaneous wound healing. ANGPTL4 deficiency impairs wound healing. H & E staining shows that ANGPTL4 deficiency significantly reduces the thickness, length and area of regenerated epidermis postwounding. Immunohistochemical staining for markers of EpSCs (α6 integrin and β1 integrin) and cell proliferation (PCNA) shows that the number and proliferation of EpSCs in the basal layer of the epidermis are reduced in ANGPTL4-deficient mice. In vitro studies show that ANGPTL4 deficiency impedes EpSC proliferation, causes cell cycle arrest at the G1 phase and reduced the expression of cyclins D1 and A2, which can be reversed by ANGPTL4 overexpression. ANGPTL4 deletion suppresses EpSC migration, which is also rescued by ANGPTL4 overexpression. Overexpression of ANGPTL4 in EpSCs accelerates cell proliferation and migration. Collectively, our results indicate that ANGPTL4 promotes EpSCs proliferation by upregulating cyclins D1 and A2 expression and accelerating cell cycle transition from G1 to S phase, and ANGPTL4 promotes skin wound re-epithelialization by stimulating EpSC proliferation and migration. Our study reveals a novel mechanism underlying EpSC activation and re-epithelialization during cutaneous wound healing.

## Introduction

Wound healing is a dynamic process involving homeostasis, inflammation, proliferation and tissue remodeling [1]. The proliferative phase is characterized by granulation tissue formation, collagen deposition, angiogenesis and re-epithelialization. Epidermal stem cells (EpSCs), located in the basal layer of the epidermis, are essential for wound repair. In response to skin injury, EpSCs proliferate, migrate and differentiate into keratinocytes to regenerate the epidermis [2]. The genes involved in regulating EpSC proliferation and migration during wound healing are not fully understood.

Angiopoietin-like 4 (ANGPTL4) is a member of the angiopoietin-like gene family. It is a multifunctional protein that has been reported to be involved in lipid metabolism, angiogenesis, stem cell regulation and various diseases, such as metabolic and cardiovascular diseases, cancer and rheumatoid arthritis [3-8]. In addition, ANGPLT4 is involved in wound healing, including wound angiogenesis [9], keratinocyte differentiation and migration [10,11] ANGPTL4 is upregulated in the epidermis at the edge of wound during wound healing [11, 12]. Knockout of the *Angptl4* gene in mice impaired keratinocyte migration and delayed wound re-epithelialization [11]. Therefore, we proposed that the elevation of ANGPTL4 postwounding might regulate EpSC proliferation and migration during wound re-epithelialization.

In this study, we investigated the involvement of ANGPTL4 in re-epithelialization of skin wound and explored the underlying mechanisms using *Angptl4* knockout (*Angptl4*^*-/-*^) mice and primary cultured murine EpSCs.

## Materials and Methods

### Animals

*Angptl4*^*-/-*^ mice obtained from the Mutant Mouse Resource and Research Center, USA, were crossed with C57BL/6 mice from the Comparative Medical Center of Yangzhou University to obtain *Angptl4*^*-/-*^ mice on the C57BL6 background. The C57BL/6 mice used in this study were also obtained from this center. *Angptl4*^*-/-*^ mice and C57BL/6 mice were bred in the same animal facility. Animals were fed a standard laboratory chow and housed in a conventional animal facility. Animals were relocated to individual cages prior to the wound healing experiments. All animal experiments were performed following the guidelines of the Animal Care and Use Committee of Suzhou Ruihua Orthopedic Hospital.

### Mouse skin wound model

Eight to ten week old wild type (WT) and *Angptl4*^*-/-*^ mice were anesthetized with pentobarbital sodium at a concentration of 25% (35 mg/kg). The animals’ backs were shaved and sterilized with iodophor. A full-thickness wound was made in the upper paravertebral region using an 8 mm biopsy punch (Haiyan Flagship Store, Suzhou, China). After wounding, the wounds were photographed every two days. The wound area was calculated using ImageJ software (NIH Image, Bethesda, Maryland, USA). Animals were sacrificed under anesthesia at 4 and 8 days after wounding, and the wounds and surrounding skin tissues were cut and fixed in formalin (10%) for further histological and immunohistochemical analyses.

### Histology and immunohistochemistry

Formalin-fixed skin tissues were embedded in paraffin, sectioned at 4 μm, and stained with hematoxylin and eosin (H & E). Primary antibodies against ANGPTL4, β1 integrin, PCNA (Abcam, Cambridge, UK) or α6 integrin (Bioworld, Texas, USA), and HRP-conjugated secondary antibodies (MXB, Fuzhou, China) were used for immunohistochemical staining. The sections were counterstained with Hematoxylin and photographed under a microscope. ImageJ software was used to analyze the thickness of the epidermis, the length of neo-epithelial tongue, the area of neo-epithelium in H& E stained sections, and the positive signals in immunohistochemically stained sections.

### Isolation, culture and characterization of mouse EpSCs

EpSCs were isolated from the skin of newborn mice as previously described [13, 14]. Briefly, skin tissues were collected from anesthetized mice, cut into 1.0×1.0 cm^2^ pieces, and incubated in Dispase II (0.25%, Sigma, St Louis, USA) for 16-18 h at 4[. The epidermis was separated, minced and digested with trypsin (0.05%) for 15 min in a humidified incubator at 37°C. The dissociated tissue was filtered through a 70 µm strainer and the cell suspension was centrifuged. The cell pellet was washed and resuspended in Keratinocyte Growth Medium-2 (KGM2, PromoCell, Heidelberg, Germany) containing bovine pituitary extract (4 μl/mL), epidermal growth factor (0.125 ng/mL), insulin (5 μg/mL), epinephrine (0.39 μg/mL), hydrocortisone (0.33 μg/mL), CaCl_2_ (0.06 mM), transferrin (10 μg/mL) and antibiotics, and seeded onto plates precoated with type IV collagen. After incubation at 37[for 10 min and rinsing with PBS. The adherent EpSCs were cultured in KGM2 supplemented with Y-27632 (10 μM), which can inhibit EpSC differentiation and promote EpSC proliferation. Further experiments were performed with the first passage of EpSCs cultured in KGM2 without Y-27632 and epidermal growth factor. Biomarkers of EpSCs were detected by immunofluorescence staining. Briefly, paraformaldehyde (4%) and Triton-X (0.5%) were used to fix and permeabilize EpSCs. The cells were incubated with goat serum (10%) for 30 min, washed, and incubated with primary antibodies against CK19 or β1 integrin (Abcam, Cambridge, UK) overnight at 4°C, and then washed with PBS and incubated with fluorescence-conjugated secondary antibodies for 1 h. After staining the nuclei with DAPI, the fluorescence signals from the cells were detected under a fluorescence microscope (Olympus, Tokyo, Japan).

### Plasmid construction and cell transfection

To construct the ANGPTL4 expression plasmid, RNA isolated from mouse EpSCs was reverse transcribed into cDNA. The cDNA of *Angptl4* was amplified by PCR and inserted into the pCD513B-1 vector (Thermo, California, USA) using the ClonExpress Ultra One Step Cloning Kit (Vazyme Biotech Co., Nanjing, China). *Angptl4* cDNA sequence was confirmed by sequencing. WT EpSCs were transfected with pCD513B-1 vector or ANGPTL4 expression plasmid, and *Angptl4*^*-/-*^ EpSCs were transfected with pCD513B-1 vector or ANGPTL4 expression plasmid using Lipofectamine 3000 (Invitrogen, Massachusetts, USA). Twenty-four hours later, the cells were split into 96-well plates and cultured for different periods of time to examine cell proliferation, or transferred to 6-well plates to perform cell migration assay. Forty-eight hours later, the cells were harvested to analyze the cell cycle phase distribution by flow cytometry analysis, and to examine the expression of cyclins by RT-PCR.

### Cell proliferation assays

MTT and BrdU incorporation assays were performed to measure the proliferation of EpSCs as previously described [13, 14]. Optical density values were measured at 490 nm for MTT assay and at 450 nm for BrdU incorporation assay, using the Multiskan™ Spectrum Microplate Reader (Thermo Fisher, Massachusetts, USA).

### Cell migration assay

The scratch-wound healing assay was performed to examine the migration of mouse EpSCs. EpSCs grown to confluence in 6-well plates were treated with mitomycin C (10 μg/mL) for 2 h and washed with PBS. The cell monolayer was scratched with a 10 μL pipette tip, washed with PBS, and cultured at 37°C. Images were taken at 0, 12 and 24 h after scratching using an inverted phase contrast microscope (Olympus, Tokyo, Japan). The area of the gap that was not covered by EpSCs was analyzed using the ImageJ software.

### Flow cytometry analysis

Flow cytometry was used to analyze the expression of EpSC biomarkers. After suspension in FACS buffer (PBS, 1% goat serum, 5% FBS) for 1 h at 25°C, mouse EpSCs were incubated with PE-labeled CD71 antibody and FITC-conjugated CD49f antibody (BD Biosciences, San Jose, USA). PE/FITC-conjugated IgG2a was used as an isotype control. Cells were washed, resuspended in FACS buffer and analyzed by flow cytometry (Beckman Coulter, California, USA).

The cell cycle was analyzed by flow cytometry. Cultured EpSCs were harvested and fixed in ethanol (75%) overnight at 4 °C. After ethanol was removed by centrifugation, cells were stained with PI/RNase Staining Buffer (BD Biosciences, San Jose, USA) for 30 min in the dark and analyzed by flow cytometry.

### Reverse transcriptase-polymerase chain reaction (RT-PCR)

The TRIzol reagent (Invitrogen, Carlsbad, USA) was used to extract total RNA from mouse EpSCs. cDNA was reverse transcribed from RNA using the HiScript II Reverse Transcriptase (Vazyme Biotech Co., Nanjing China). The PCR reaction was performed with 40 cycles of 95°C for 30 seconds, 55 [for 40 seconds, and 72 [for 1 min. PCR products were identified by agarose gel electrophoresis and ethidium bromide staining. Semiquantitative analysis of target gene expression was performed using the ImageJ software. The PCR primer sequences (5′-3′) are as follows: *Angptl4*: GGTGAATAAGAGGAGGTTGC (forward), CCGATTGTCTGTTGTGCC (reverse); *Gapdh*: GCAGTTCGC CTTCACTATGGA (forward), ATCTTGTGGCTTGTCCTCAGAC (reverse); Cyclin D1: GTGCGTGCAGAAGGAGATTGT (forward), CAGCGGGAAGACCTCCTCTT (reverse); Cyclin E1: TCTCCTCACTGGAGTTGATGCA (forward), AACGGAACCATCCATTTGACA (reverse); cyclin A2: CCTTCCACTTGGCTCTCTACACA (forward), GACTCTCCAGGGTATATCCAGTCTGT (reverse).

### Western blot

EpSCs were lysed in RIPA lysis buffer. After centrifugation, the supernatant was collected and the protein concentration was measured using a Bradford protein assay kit (Beyotime, Wuhan, China). Western blot was performed according to general protocols. Primary antibodies against ANGPTL4, cyclin A2, cyclin D1 and GAPDH (Abcam, Cambridge, UK) were used. The target proteins were visualized using the High-Sensitivity ECL Chemiluminescence Detection Kit (Vazyme Giotech Co., Nanjing China), and quantified using ImageJ software.

### Statistical analysis

All results are expressed as mean ± standard deviation (SD). A two-tailed Student’s t-test was used to analyze the difference between two groups. A P-value equal to or less than 0.05 was considered to be statistically significant.

## Results

### ANGPTL4 deficiency delays skin wound healing

To investigate the contribution of ANGPTL4 to cutaneous wound repair, the expression of ANGPTL4 was examined in uninjured skin tissue and in the skin tissue adjacent to the full-thickness wounds on days 4 and 8 after injury in WT mice. Immunohistochemical staining of normal skin sections showed that ANGPTL4 was mainly expressed in the epidermis and hair follicles. ANGPTL4 was upregulated in the basal layer cells of the regenerated epidermis during the wound healing process (Figure 1A). Knockout of *Angptl4* greatly reduced its expression in the epidermis (Figure 1B). The effect of ANGPTL4 on skin wound repair was examined using *Angptl4* knockout mice. Figure 1C shows the reduction in wound area over time and demonstrates that wound closure was slower in *Angptl4*^*-/-*^ mice than in WT mice. Taken together, these data indicate that ANGPTL4, which is expressed in the basal layer cells of the epidermis, contributes significantly to cutaneous wound healing.

**Figure 1.**
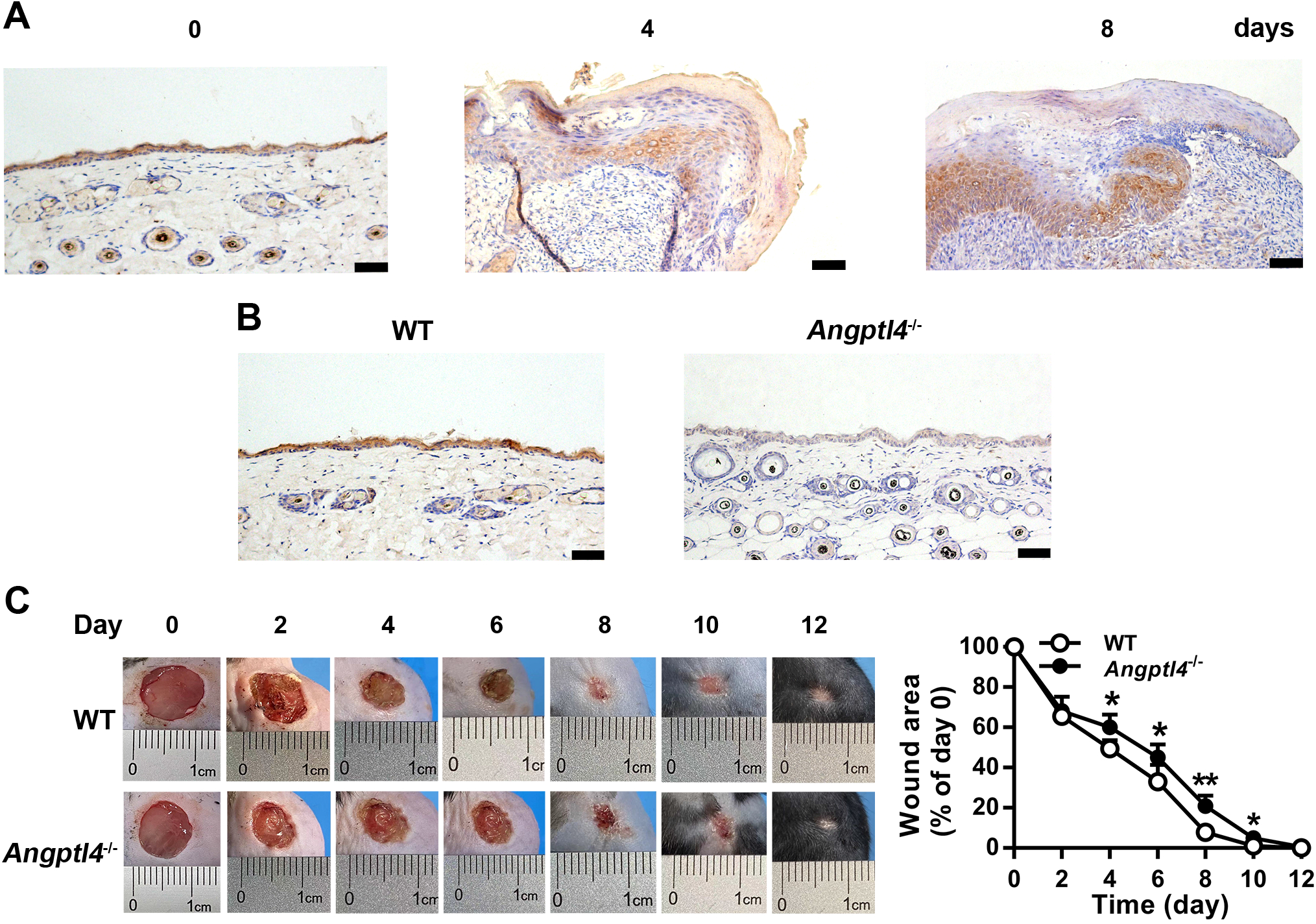
ANGPTL4 deficiency delays cutaneous wound healing in mice. (A) Representative images of immunohistochemical staining for ANGPTL4 in normal skin tissue, and in skin tissue at the wound edge during wound healing. (B) Detection of ANGPTL4 expression in skin tissues of WT and *Angptl4*^*-/-*^ mice by immunohistochemical staining. (C) Representative photographs of skin wound healing over time in WT and *Angptl4*^*-/-*^ mice, and changes in wound area measured by ImageJ. Data are mean ± SD, n=6/group. \**P* < 0.05, \*\**P* < 0.01, compared with WT mice. Scale bars are 50 μm in (A) and (B).

### ANGPTL4 deficiency impairs wound re-epithelialization

To investigate whether ANGPTL4 affects the regeneration of the epidermis after wounding, we examined the histology of uninjured skin tissues and skin tissues adjacent to the wounds on days 4 and 8 after injury in WT and *Angptl4*^*-/-*^ mice. H & E staining showed that *Angptl4* deletion had no significant effect on the cell distribution in the epidermis and the thickness of the epidermis (Figure 2A and B). However, the thickness of the regenerated epidermis in *Angptl4*^*-/-*^ mice was smaller than that in WT mice (Figure 2A and B). The length of the regenerated epidermis (neo-epithelial tongue) (Figure 2A and C) and the area of the regenerated epidermis (neo-epithelium) (Figure 2A and D) were also significantly reduced in *Angptl4*^*-/-*^ mice. These data demonstrate that ANGPTL4 plays a crucial role in the re-epithelialization of cutaneous wounds.

**Figure 2.**
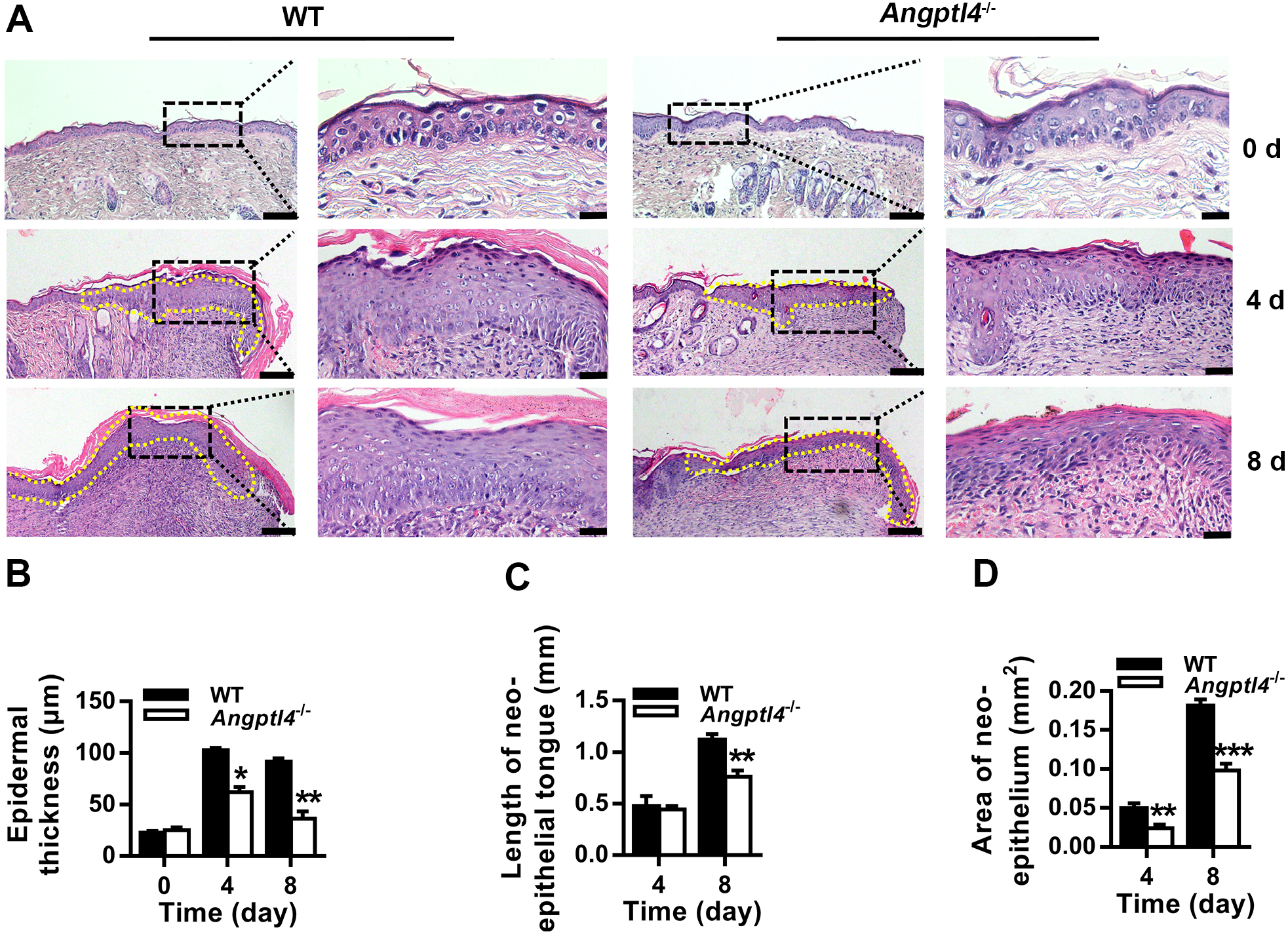
ANGPTL4 deficiency impairs wound re-epithelialization. (A) Representative H & E staining images of uninjured skin tissue from WT and *Angptl4*^*-/-*^ mice, and of regenerated skin tissue adjacent to wounds on days 4 and 8 after injury (A). In each group, the scale bars are 100 μm and 20 μm in the left and right panels, respectively. (B-D) Epidermal thickness (B), neoepithelial tongue length (C) and neoepithelial area were measured using ImageJ (D). Data are mean ± SD, n =6/group. \**P* < 0.05, \*\**P* < 0.01, \*\*\**P* < 0.001, compared with WT mice.

### ANGPTL4 deficiency inhibits EpSC proliferation in the epidermis adjacent to the wound

To investigate whether ANGPTL4 promotes wound healing by promoting the proliferation of EpSCs in the epidermis, the expression of biomarkers of cell proliferation (PCNA) and EpSCs (β1 integrin and α6 integrin) were detected by immunohistochemical staining. As shown in Figure 3, the cells expressing PCNA, β1 integrin and α6 integrin were located in the basal layer of the epidermis. The number of these cells in uninjured skin was not different between WT and *Angptl4*^*-/-*^ mice. PCNA-, β1 integrin- and α6 integrin-expressing cells were greatly increased in the regenerated epidermis of both WT and *Angptl4*^*-/-*^ mice on days 4 and 8 after wounding, but the number of these cells was lower in *Angptl4*^*-/-*^ mice than in WT mice. Because ANGPTL4 was upregulated in the basal layer cells of the regenerating epidermis after skin injury (Figure 1A), the above results indicate that ANGPTL4 stimulates EpSC proliferation during wound healing.

**Figure 3.**
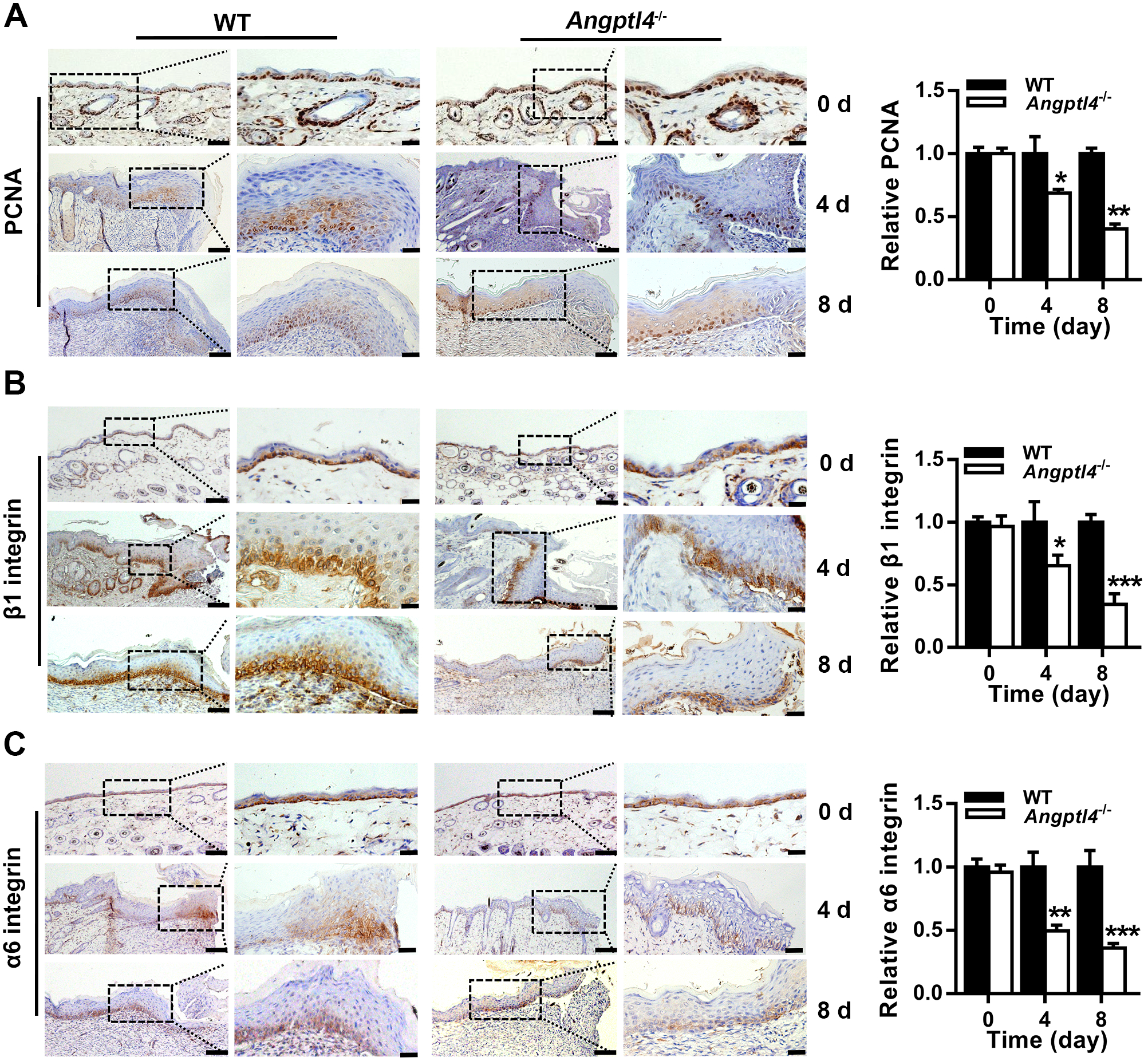
ANGPTL4 deficiency hinders the proliferation of EpSCs in skin tissues during wound healing. Representative images of immunohistochemical staining for PCNA (A), β1 integrin (B), and α6 integrin (C) in uninjured skin tissues, and of wound edge skin tissues on days 4 and 8 after injury from WT and *Angptl4*^*-/-*^ mice. Data are mean ± SD, n=3/group. \**P* < 0.05, \*\**P* < 0.01, \*\*\**P* < 0.001, compared with WT mice. In each group, the scale bars are 100 μm and 20 μm in the left and right panels, respectively.

### ANGPTL4 promotes EpSC proliferation

We isolated EpSCs from murine skin tissues to examine the effect of ANGPTL4 on EpSC proliferation and migration. The cultured EpSCs showed cobblestone-like morphology under a light microscopy (Figure 4A). Flow cytometry assays were performed to examine the expression of α6 integrin and CD71 (Figure 4B), two commonly-recognized markers of EpSC, in these cells. The population of α_6_ integrin^high^/CD71^low^ cells was 98% (Figure 4B). Immunofluorescence staining showed high expression of β_1_ integrin and CK19, two other markers of EpSCs, by these cells. These results demonstrated the high-purity isolation of murine skin EpSCs. MTT assay showed that the proliferation of ANGPTL4-deficient EpSCs was slower than that of WT EpSCs (Figure 5A). BrdU incorporation assay confirmed the impairment of proliferation caused by ANGPTL4 deletion (Figure 5B). Transfection of ANGPTL4-deficient EpSCs with ANGPTL4 expression plasmid increased ANGPTL4 levels and restored cell proliferation (Figure 5C and D). Overexpression of ANGPTL4 in WT EpSCs enhanced cell proliferation (Figure 5E and F), demonstrating the proproliferative effect of ANGPTL4 on EpSCs.

**Figure 4.**
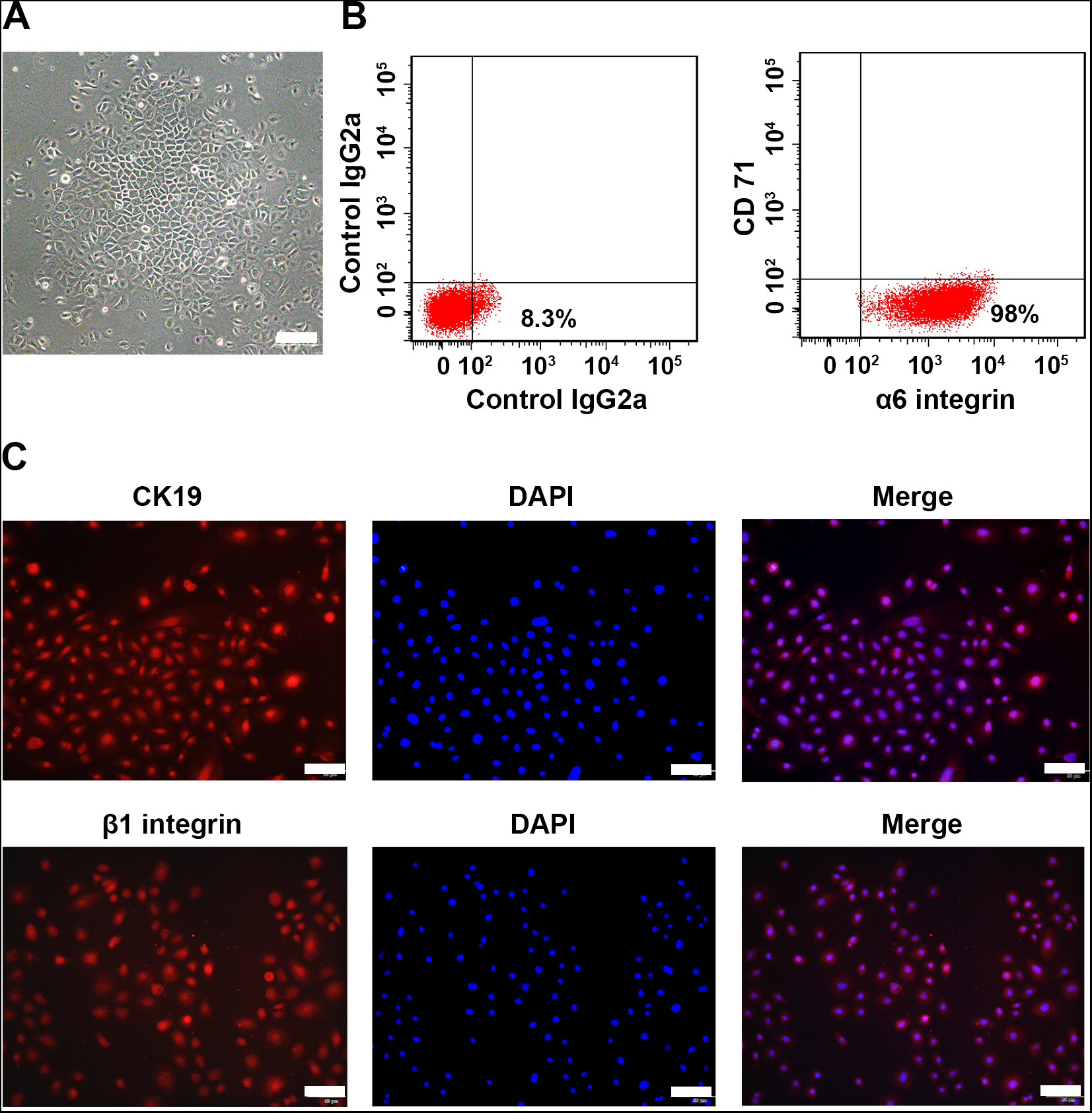
Characterization of mouse EpSCs. (A) Morphology of cultured mouse EpSCs under a light microscopy. Scale bar = 100 μm. (B-C) Expression of EpSCs biomarkers α6 integrin, CD71, β1 integrin and CK19 in cultured EpSCs was examined by flow cytometry analysis (B) and immunofluorescence staining (C). Scale bar = 50 μm.

**Figure 5.**
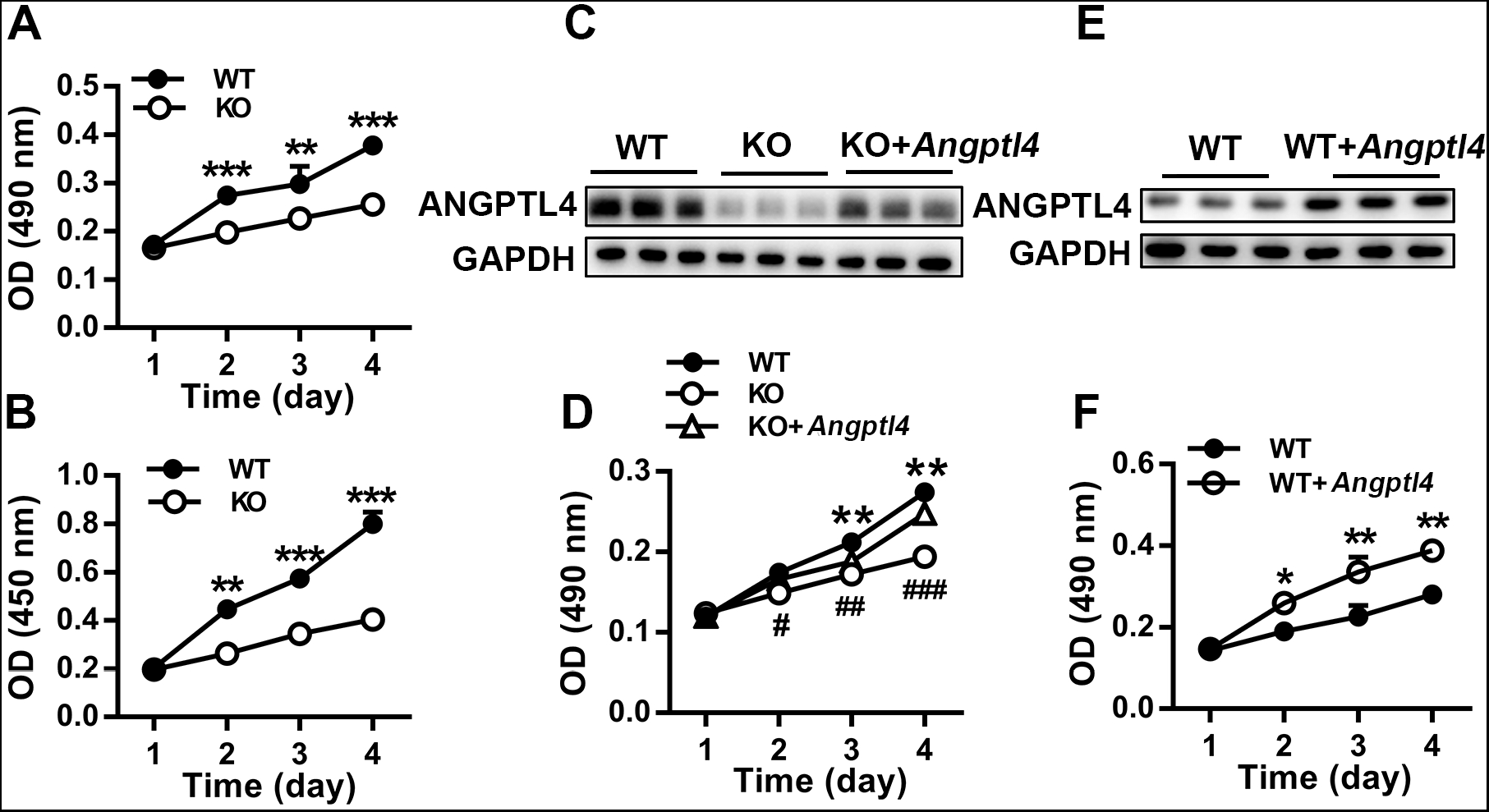
ANGPTL4 promotes EpSC proliferation. (A-B) Proliferation of EpSCs isolated from WT and *Angptl4*^*-/-*^ mice was determined by MTT assay (A) and BrdU incorporation assay (B). (C-F) WT EpSCs transfected with control plasmid (WT) and *Angptl4*^*-/-*^ EpSCs transfected with control plasmid (KO) or ANGPTL4 expression plasmid (KO+*Angptl4*) (C, D); WT EpSCs transfected with control plasmid (WT) or ANGPTL4 expression plasmid (WT+*Angptl4*) (E, F) were analyzed for ANGPTL4 expression by Western blot (C, E) and for cell proliferation by MTT assay (D, F), respectively. Data are mean ± SD, n=3. \**P* < 0.05, \*\**P* < 0.01, \*\*\**P* < 0.001, comparison between WT and KO EpSCs (A, B, D) or between WT and (WT+*Angptl4*) EpSCs (F); ^#^*P* < 0.05, ^##^*P* < 0.01, ^###^*P*<0.001, comparison between (KO+*Angptl4*) and KO EpSCs.

### ANGPTL4 regulates cell cycle of EpSCs and cyclins expression

The effect of ANGPTL4 on the cell cycle distribution of EpSCs was examined by flow cytometry analysis. As shown in Figure 6A, ANGPTL4 deletion in EpSCs significantly increased the cell population in G1 phase, and decreased the proportion of cells in G2/M phase. Transfection of ANGPTL4-deficient EpSCs with ANGPTL4 expression plasmid significantly decreased the number of cells in G1 phase and increased the number of cells in S phase. These results indicate that ANGPTL4 deficiency suppresses EpSC proliferation by inducing cell cycle arrest in G1 phase, and overexpression of ANGPTL4 stimulates EpSC proliferation by promoting G1 to S phase transition. Since cyclins play an important role in cell cycle progression, the effect of ANGPTL4 on cyclins D1 and A2 expression was examined at both mRNA and protein levels. Deletion of ANGPTL4 in EpSCs significantly decreased the expression of cyclins D1 and A2 expression. Overexpression of ANGPTL4 in ANGPTL4-deficient EpSCs upregulated the expression of these molecules (Figure 6B and C). These data indicate that ANGPTL4 stimulates EpSC proliferation by inducing the expression of cyclins D1 and A2.

**Figure 6.**
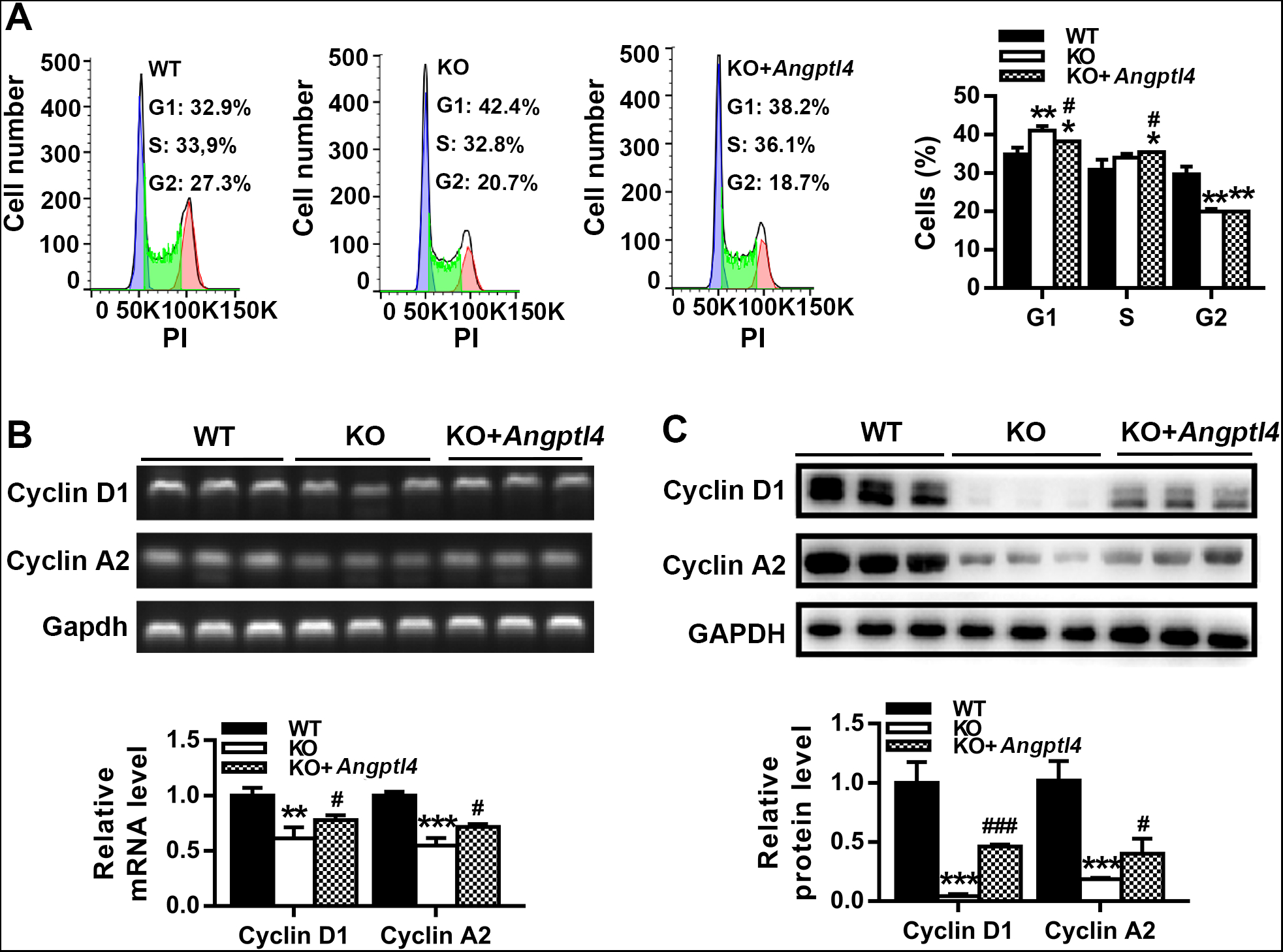
ANGPTL4 regulates cell cycle and cyclins expression in EpSCs. WT EpSCs transfected with control plasmid (WT), and *Angptl4*^*-/-*^ EpSCs transfected with control plasmid (KO) or ANGPTL4 expression plasmid (KO+*Angptl4*) were examined for cell cycle distribution by flow cytometry analysis (A) and for cyclins D1 and A2 expression at mRNA and protein levels (B, C), respectively. Data are mean ± SD, n=3. \**P* < 0.05, \*\**P* < 0.01, \*\*\**P* < 0.001, compared with WT EpSCs; ^#^*P* < 0.05, ^###^*P* < 0.001, compared with KO EpSCs.

### ANGPTL4 enhances EpSC migration

The effect of ANGPTL4 on the migration of EpSCs was examined using the scratch-wound healing assay in vitro. As shown in Figure 7A, ANGPTL4 deletion significantly suppressed the migration of EpSCs and delayed the closure of the scratch wound. Overexpression of ANGPTL4 in ANGPTL4-deficient EpSCs restored cell migration. Furthermore, overexpression of Angptl4 in EpSCs promoted cell migration (Figure 7B).These results indicate that ANGPTL4 contributes to EpSC migration and elevation of Antgptl4 accelerates EpSC migration.

**Figure 7.**
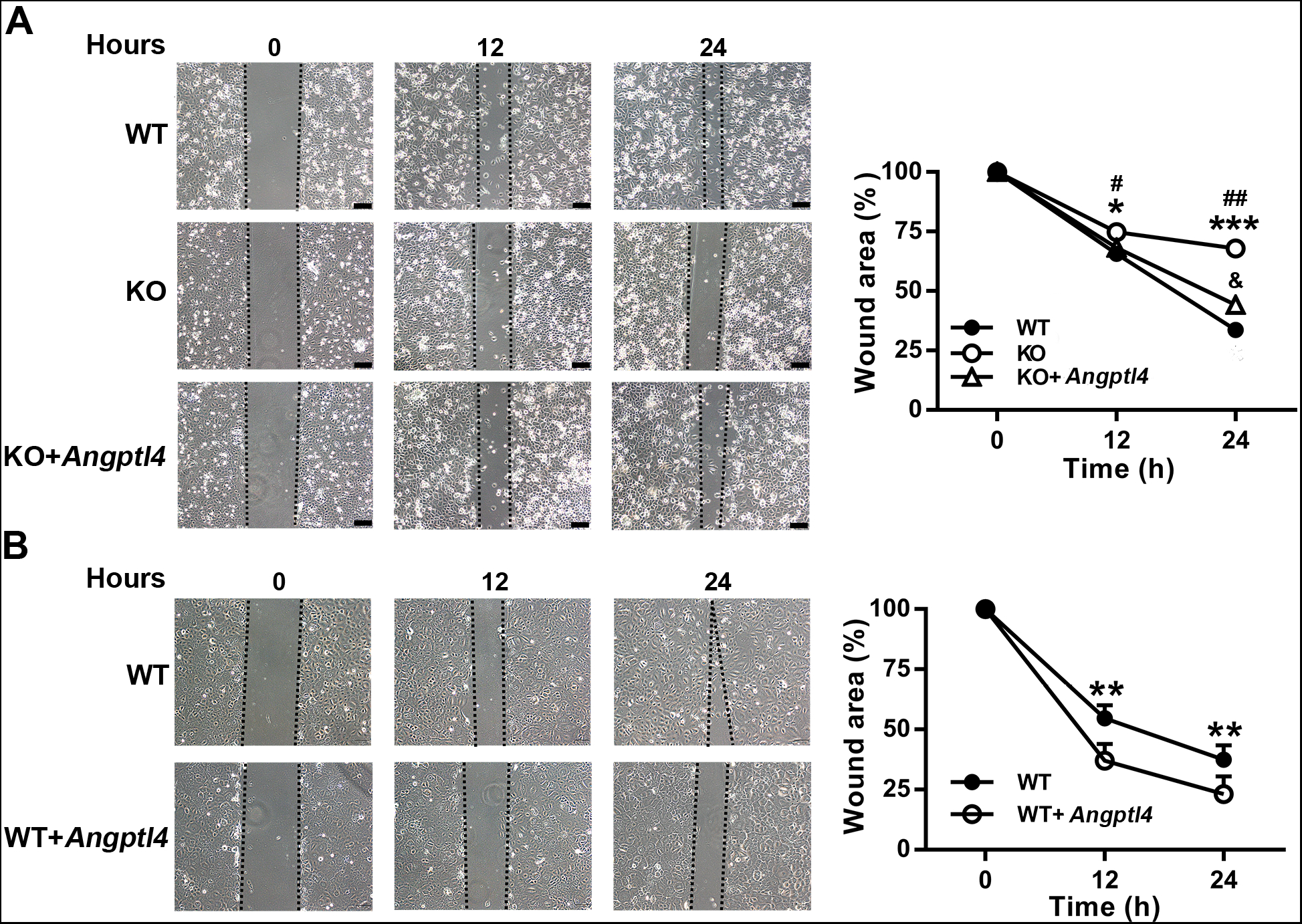
ANGPTL4 promotes EpSC migration. Wild type EpSCs transfected with control plasmid (WT) and *Angptl4*^*-/-*^ EpSCs transfected with control plasmid (KO) or ANGPTL4 expression plasmid (KO+*Angptl4*) (A); wild type EpSCs transfected with control plasmid (WT) or ANGPTL4 expression plasmid (WT+*Angptl4*) (B) were examined for cell migration by scratch wound healing assay. Data are mean ± SD, n=3. A: \**P* < 0.05, \*\*\**P* < 0.001, comparison between KO EpSCs and WT EpSCs; ^#^*P* < 0.05, ^##^*P* < 0.01, comparison between KO EpSCs and (KO+*Angptl4*) EpSCs; ^&^P<0.05, comparison between (KO+*Angptl4*) EpSCs and WT EpSCs. B: \*\**P* < 0.01, comparison between WT and (WT+*Angptl4*) EpSCs. Scale bar = 100 μm.

## Discussion

ANGPTL4 has been reported to play an important role in skin wound repair by regulating monocyte-to-macrophage differentiation during the inflammatory phase [15], keratinocyte migration and differentiation during wound re-epithelialization [10,11,16]. Topical application of recombinant ANGPTL4 accelerates wound healing in diabetic mice by improving angiogenesis [9]. In this study, we found that ANGPTL4 contributed to cutaneous wound re-epithelialization by stimulating the proliferation and migration of EpSCs.

The balance between self-renewal and differentiation of EpSCs plays a key role in the homeostasis of the skin epidermis. The skin epidermis is regenerated by EpSCs proliferation and migration in response to skin injury. We found that knockout of *Angptl4* in mice had no significant effect on epidermal thickness and the number of EpSCs in the epidermis (Figure 2A, Figure 3), indicating that ANGPTL4 is not involved in epidermal homeostasis. Our study showed that ANGPTL4 expression was upregulated in the basal layer cells of the epidermis adjacent to the wound during wound healing (Figure 1A). Deficiency of ANGPTL4 in mice impaired EpSC proliferation (Figure 3), wound re-epithelialization (Figure 2), and wound closure (Figure 1C) after skin injury. These results strongly suggest that elevation of ANGPTL4 promotes EpSC proliferation during wound healing. The factors that stimulate *Angptl4* expression during wound healing are not clear. Singh et al. [17] recently reported that platelet-derived growth factors can stimulate *Angptl4* expression in cultured keratinocytes and skin explants. Since platelets are activated and release various active molecules after skin injury [18], we speculate that the growth factors released by the activated platelets may contribute to the upregulation of ANGPTL4 in the epidermis.

To investigate whether ANGPTL4 could directly stimulate EpSC proliferation, we isolated and cultured EpSCs from WT and *Angptl4*^*-/-*^ mice. In vitro study showed that ANGPTL4 deficiency impaired EpSC proliferation (Figure 5A and B). Rescuing ANGPTL4 expression in *Angptl4*^*-/-*^ EpSCs by transfecting the cells with Angptl4 expression plasmid significantly enhanced cell proliferation (Figure 5D). Elevation of ANGPTL4 in WT EpSCs by transfecting the cells with ANGPTL4 expression plasmid significantly stimulated cell proliferation (Figure 5E and F). These results demonstrate that the basal level of ANGPTL4 is essential for EpSC proliferation and that increasing ANGPTL4 promotes EpSC proliferation. To gain further insight into the underlying mechanisms of ANGPTL4-regulated EpSC proliferation, we determined the cell cycle distribution of WT and ANGPTL4-deficient EpSCs. The results showed that ANGPTL4 deficiency caused cell cycle arrest in the G1 phase, and reduced the cell population in the G2/M phase. Overexpression of ANGPTL4 promoted cell cycle progression from G1 to S phase (Figure 6A). Cyclins control cell cycle progression by activating cyclin-dependent kinase (CDK). To understand the regulation of cell cycle progression by ANGPTL4, we examined the expression of cyclins involved in the regulation of G1 phase progression and the transition from G1 to S phase. Cyclin D1 controls the G1 to S phase transition by binding to CDK4/CDK6. Cyclin A activates CDK1 and CDK2 and plays a critical role in DNA replication during the S phase, and also functions in the G2-to M-phase transition [19]. Our study showed that ANGPTL4 deficiency in EpSCs decreased the expression of cyclins D1 and A2, whereas ANGPTL4 overexpression upregulated the expression of these molecules. These results indicate that ANGPTL4 stimulates the proliferation of EpSCs by upregulating the expression of cyclins D1 and A2. The signaling pathway mediating the regulation of cyclins D1 and A2 by ANGPTL4 requires further investigation.

Our in vitro studies showed that ANGPTL4 deficiency impaired EpSC migration, which could be reversed by ANGPTL4 overexpression (Figure 7A). Overexpression of ANGPTL4 in EpSCs promoted cell migration (Figure 7B). These results indicate that the basal level of ANGPTL4 plays an important role in EpSC migration, and that elevating ANGPTL4 accelerates EpSC migration. Goh et al. [11] reported that inhibition of ANGPTL4 in human keratinocytes by a neutralizing antibody or siRNA impaired cell migration. ANGPTL4 modulates keratinocyte migration by interacting with integrins β1 and β5 and activating the FAK-Src-PAK1 signaling pathway. We speculate that the signaling pathway by which ANGPTL4 stimulates EpSC and keratinocyte migration may be similar, but needs to be verified.

In conclusion, ANGPTL4 promotes EpSC proliferation by upregulating cyclin A2 and D1 expression and accelerating the cell cycle transition from G1 to S phase. Elevation of ANGPTL4 in the epidermis adjacent to the wound after skin injury may contribute to re-epithelialization and wound healing by promoting proliferation and migration of EpSCs. ANGPTL4 is a potential therapeutic agent for the treatment of poorly-healing wounds.

